# Operative temperatures of eastern garter snakes (*Thamnophis sirtalis sirtalis*) reveal a Goldilocks effect for habitat use

**DOI:** 10.1101/2025.06.11.659115

**Authors:** Harry A. W. Kumbhani, Curtis R. Abney, Danilo Giacometti, Glenn J. Tattersall

## Abstract

Garter snakes (*Thamnophis* spp.) are the most widespread reptiles in North America, although evidence suggests that thermal preference has not diverged much among populations or *Thamnophis* species. To shed light on how thermal decisions influence local habitat use by the eastern garter snake (*Thamnophis sirtalis sirtalis*), we measured the thermal profiles of three habitats differing in canopy cover: open peat, mixed shrub, and closed forest. We installed biophysical models to record operative temperatures at a fine scale and assess habitat thermal quality. We also used coverboards to survey habitat usage. While the open canopy offered the highest thermal quality, we recorded the greatest number of snakes in the mixed shrub which had a lower thermal quality. Since environmental temperatures regularly exceeded the upper thermal limit of *T. s. sirtalis* in the open canopy, snakes might favour the use of habitats that minimise the odds of overheating. Therefore, open habitats potentially restrict snakes’ activity window and may not be thermally attractive. Our data show that *T. s. sirtalis* use habitats that vary in thermal quality, but warmer habitats are not necessarily better. Rather, snakes preferentially seek areas that offer a mix of open and closed canopies to suit their thermoregulatory needs.

## Introduction

Ectotherms are thought to exhibit an innate preference for a certain range of body temperatures (Huey and Slatkin 1976; Hertz et al. 1993). Preferred body temperatures (T_set_) represent the fundamental thermal niche (Magnuson et al. 1979), and are argued to encompass the optimal body temperatures for physiological processes such as growth, development, reproduction, maintenance, digestion, and physical performance (Huey and Slatkin 1976; Huey 1991; Hertz et al. 1993; Ming-Chung Tu and Hutchison 1994; Row and Blouin-Demers 2006a; Lelièvre et al. 2011). Consequently, individuals are presumed to strive to maintain T_set_ when active in the wild and when environmental conditions are favourable (Hertz et al. 1993; Row and Blouin-Demers 2006a; Martin and Huey 2008). However, T_set_ assessments are primarily performed in laboratory settings, where thermoregulatory costs are minimised, meaning that the extent to which animals are capable of realising T_set_ in biologically relevant scenarios is often unclear (Huey 1991; Hertz et al. 1993; Row and Blouin-Demers 2006a; Giacometti et al. 2021).

The T_set_ of many reptiles lies close to their upper temperature threshold (CT_max_), above which physiological function is impaired (Huey and Slatkin 1976; Martin and Huey 2008; Cadena and Tattersall 2009; Gilbert and Miles 2017; Huey and Pianka 2017). Reptiles can remain active at a wide range of temperatures below their T_set_, but only a narrow range above their T_set_ (Martin and Huey 2008; Huey and Pianka 2017). Despite reptiles experiencing reduced performance when exposed to temperatures below their T_set_ range, being too cool is rarely fatal (Spellerberg 1972; Doughty 1994; Martin and Huey 2008). Therefore, animals may prefer to maintain slightly lower body temperatures (T_b_) than to operate in hotter thermal environments (Cadena and Tattersall 2009; Huey and Pianka 2017). In the wild, reptiles may maintain their body temperatures at the lower end of their T_set_ range thereby sacrificing their maximum performance to avoid overheating (Martin and Huey 2008; Huey and Pianka 2017). Indeed, temperate reptiles may occupy habitats where their T_set_ is rarely available and thus often operate below their T_set_ (Blouin-Demers and Weatherhead 2001; Martin and Huey 2008).

For reptiles the possibility to achieve T_set_ is an integral part of habitat selection (Gifford et al. 2012; Halliday and Blouin-Demers 2016; Lelièvre et al. 2011; Row and Blouin-Demers 2006a, 2006b). Thermal quality (*d*_e_) is a way of measuring the quality of a habitat available based on the thermal needs of an animal (Hertz et al. 1993). This metric is defined as the absolute deviation between operative environmental temperature (T_e_) and a species’ T_set_ (Hertz et al. 1993). Thus, *d*_e_ provides an estimate of the thermoregulatory effort required to achieve T_set_ within a given habitat. Lower *d*_e_ values indicate the target species is close to its T_set_, hypothetically paying lower thermoregulatory costs to achieve this temperature (Hertz et al. 1993). By contrast, higher *d*_e_ values indicate the target species is far from its T_set_, and would likely pay greater thermoregulatory costs to achieve this temperature (Hertz et al. 1993). Therefore, low *d*_e_ values indicate a high thermal quality habitat while high *d*_e_ values indicate a low thermal quality habitat for the target-species (Hertz et al. 1993). In a heterogenous terrestrial environment, reptiles can choose between different substrates above ground (e.g. arboreal species), below ground (e.g. fossorial species), or on the ground itself (Blouin-Demers and Weatherhead 2001; Brown and Weatherhead 2000; Lelièvre et al. 2011; Row and Blouin-Demers 2006b). Furthermore, gradients of sun exposure may exist within an environment, creating habitats with different amounts of sunlight (Row and Blouin-Demers 2006b; Elmore et al. 2017). Several habitat properties may impact *d*_e_, including the level of solar radiation, air temperature, substrate temperature, and habitat humidity (Bakken 1992). As these environmental properties interact with each other, *d*_e_ may vary dramatically across small spatial scales in a heterogenous landscape (Sears et al. 2016), with important implications for thermoregulation (Giacometti et al. 2023).

In temperate snakes, *d*_e_ has been argued to be an accurate predictor of habitat quality (Halliday and Blouin-Demers 2016; Row and Blouin-Demers 2006a, 2006b). For example, eastern milk snakes (*Lampropeltis triangulum triangulum*) at the northern extent of their range preferred open habitats over forest habitats. At these more extreme latitudes, heavily shaded habitats are cooler, have high *d*_e_ values, and thus offer fewer thermoregulatory opportunities to surface-dwelling snakes. As such, the poor thermal quality of forests leads to forest-dwelling snakes expending more energy to thermoregulate. By contrast, open habitats receive more solar radiation, which translates to higher temperatures during the day, lower *d*_e_ values, and more thermoregulatory opportunities (Row and Blouin-Demers 2006b). Basking areas are also limited in forest habitats and may, with consistent use over time, become predictable locations for predation (Row and Blouin-Demers 2006a). The thermoregulatory benefit of open areas is further underscored by the fact that *L. t. triangulum* were found in greater numbers in the open habitat than the forest habitat despite their prey being more abundant in the forest habitat (Row and Blouin-Demers 2006b). The common garter snake (*Thamnophis sirtalis*) was also found to prefer open habitats over forest habitats regardless of food and conspecific density (Halliday & Blouin-Demers, 2016). Indeed, various studies have recurrently noted *T. sirtalis*’s affinity for open habitats, and thermal benefit has been argued to be one of the explanations behind this pattern (Charland and Gregory 1995; Engelstoft and Ovaska 2000; Kjoss and Litvaitis 2001; Halliday and Blouin-Demers 2016; Diaz and Blouin-Demers 2017). In temperate regions, closed-forest canopies buffer against intense solar radiation and heat dissipation causing more stable temperature conditions with less extreme temperature fluctuations. These more stable but lower temperatures may limit *T. sirtalis*’s access to suitable temperatures (Charland and Gregory 1995; Halliday and Blouin-Demers 2016; Diaz and Blouin-Demers 2017) and therefore may constrain behavioural thermoregulation. Together, these pieces of evidence suggest that temperate snakes at northern latitudes find temperature a more limiting resource than food or interactions with conspecifics (Huey and Slatkin 1976; Row and Blouin-Demers 2006a, 2006b; Halliday and Blouin-Demers 2016).

Since air temperature alone is not an accurate indicator of a reptile’s experienced temperature in the field due to solar heating or convective cooling from wind, T_e_ is often used to characterise environmental temperatures at the organismal level (Bakken 1992). Measurements of T_e_ represent the body temperature of an organism that is in thermal equilibrium with its immediate environment (i.e. non-thermoregulating), subjected to the prevailing convective, radiative, and conductive heat transfer conditions (Hertz et al. 1993; Thompson et al. 2018). By implanting temperature loggers inside reptile-mimicking models that match the morphological characteristics and thermal inertia of a living animal, one can obtain species-specific T_e_ measurements reflective of the natural thermal environment (Hertz et al. 1993; Shine and Kearney 2001; Dzialowski 2005). Such models capture the real time temperatures a hypothetical reptile would experience while accounting for the influence of abiotic factors such as wind, solar radiation, and substrate heat conduction (Hertz et al. 1993; Vitt and Sartorius 1999; Shine and Kearney 2001; Dzialowski 2005), allowing one to accurately predict species-specific body temperatures within a specific environment assuming absence of active thermoregulation (i.e., a null model of thermoregulation).

In this study, we assessed the thermal profiles of interconnected closed-forest, mixed-shrub, and open-peat habitats that make up a landscape inhabited by the eastern garter snake (*T. s. sirtalis*). Specifically, we deployed snake analogues to measure T_e_ across habitats and calculated *d*_e_ based on population-specific T_set_ data (Giacometti et al. 2021). This allowed us to determine whether different habitats offered different thermoregulatory opportunities for *T*. *s*. *sirtalis*. We also surveyed each habitat to quantify snake abundance with the goal of assessing whether patterns of habitat use were associated with habitat quality. We hypothesised that *d*_e_ would differ among habitats, and that *T. s. sirtalis* would be more abundant in the most thermally favourable habitat. By combining these approaches, we aim to understand how thermoregulatory opportunities contribute to habitat selection decisions in temperate snakes.

## Materials and Methods

### Study species

Garter snakes (*Thamnophis* spp.) are among the most common and widespread reptiles in North America and have become so widespread due to their ability to live in a variety of habitats. Despite their wide range of habitats, T_set_ has not significantly diverged among populations of *Thamnophis* spp. (Rosen 1991). Flexible thermoregulatory behaviours and habitat use may facilitate the large range of thermal habitats garter snakes occupy. The common garter snake (*T*. *sirtalis*) has the largest range and highest abundance of all North American reptile species (Larsen 1987; Larsen et al. 1993; Shine et al. 2001; Krause et al. 2003). *Thamnophis sirtalis* possess many subspecies with morphological and ecological differences. The eastern garter snake (*T. s. sirtalis*) is one of the most widespread of these subspecies inhabiting much of eastern North America. *Thamnophis. s. sirtalis* is a medium-sized snake with adult male snout-vent length ranging between 44-60 cm and females between 55-91 cm (King 1989; Larsen et al. 1993; Krause et al. 2003).

### Study site habitat characteristics

The study site is a wetland complex in the Niagara region of southern Ontario, Canada within the Carolinian forest ecoregion of North America. Certain areas within the wetland were previously mined for peat extraction in the 20^th^ century (Yagi and Litzgus 2012). Despite this recent disturbance there is a high diversity of wetland habitats, including deciduous swamps, peatlands, and bogs (Yagi and Litzgus 2012), providing a gradation in canopy cover and thermal environments. Open habitats in temperate regions like those found in southern Ontario provide superior thermal quality for North American snake species compared to forested habitats (Blouin Demers and Weatherhead 2002; Row and Blouin-Demers 2006b; Halliday and Blouin-Demers 2016). However, little is known about intermediate successional areas which may exhibit mixed characteristics of both open and shaded environments. Inclusion of mixed shrub canopy may also provide a more realistic reflection of a habitat matrix since snakes are not confined to strictly open and closed-canopy systems. Our study site provided access to a unique open habitat. The open sites from previous studies have typically been open fields or artificially cleared openings in the landscape with little to no canopy coverage (Row and Blouin-Demers 2006b; Halliday and Blouin-Demers 2016). The open habitat at our field site was distinct in that it was part of a drained wetland complex with dark, exposed peat soils sparsely vegetated by bog species like bog blueberry (*Vaccinium uliginosum*), cottongrass (*Eriophorum angustifolium*) and sundew plants (*Drosera sp.*).

Three habitat types were examined in this study: closed forest, mixed shrub, and open peat. These habitats were pre-selected and coarsely defined by abiotic features that would primarily influence the thermal properties of the habitat. Therefore, habitat characterization was focused mainly on canopy cover and how solar radiation, soil moisture, and soil temperature differed among these sites (Table 1). Canopy cover was measured 3 times a year, after foliage had emerged, while solar radiation was measured every other survey (5 times a year). Soil moisture and soil temperature were intended to be collected during every survey, but machine malfunctions resulted in soil data missing from certain surveys. Open peat habitat was slowly regenerating after being drained and harvested for peat-mining in the mid to late 20^th^ century, and exhibited an absence of contiguous forest canopy and presence of dark hummocky soils. The forest was not an historically mined area and contained mature deciduous trees providing a solid canopy cover and thick understory. The mixed shrub site served as an intermediate between the other two habitats, where canopy overstory and understory were moderate. The plant communities within the mixed shrub were in succession from previous disturbances. Canopy cover is greatest (solar radiation lowest) in the closed forest habitat, intermediate in the mixed shrub habitat, and lowest (solar radiation greatest) in the open peat habitat (Table 1), and for this reason, we refer to the different habitats based on their different canopy types.

**Table 1:** Summary of five abiotic variables (mean ± SD) measured in each of the three habitats. *N* refers to the number of surveys which measurements were taken. Measurements of an individual survey are comprised of separate days for each canopy within a week of each other. Soil measurements were taken but are incomplete due to machine malfunctions in the field.

|  | **Closed**<br>**(N=20)** | **Mixed**<br>**(N=20)** | **Open**<br>**(N=20)** |
| --- | --- | --- | --- |
| **Solar Radiation (W/m<sup>2</sup>)** | 153 ± 128 | 278 ± 235 | 437 ± 150 |
| **Relative Humidity (%)** | 61.4 ± 16.6 | 59.6 ± 19.2 | 62.1 ± 13.7 |
| **Soil Moisture (%)** | 25.9 ± 21.3 | 19.7 ± 15.6 | 27.5 ± 17.2 |
| **Soil Temperature (°C)** | 20.5 ± 3.05 | 21.0 ± 2.86 | 25.3 ± 5.89 |
| **Air Temperature (°C)** | 24.9 ± 3.78 | 25.3 ± 3.58 | 26.7 ± 5.02 |

### Habitat thermal characteristics

Black globe temperature (T_bg_) was measured to assess the combined impact of solar radiation, wind speed, and temperature within the three habitat types. Each black globe consisted of a copper sphere painted matte black with an iButton (Thermochron model DS1922L, Maxim Integrated™) suspended inside. The copper spheres were raised off the ground by 2 m. An additional iButton (Hygrochron model DS1923, Maxim Integrated™) was affixed to the pole holding the black globe, except that this iButton was held within a high reflective housing to capture an above ground relative humidity (RH) and air temperature (T_a_) measurement in each habitat (Figure S1). T_bg_, T_a_, and RH were recorded every 20 minutes continuously from May 31^st^ to September 28^th^, 2017, and May 11^th^ to September 29^th^, 2018, with brief interruptions to download data. For analysis purposes we focused on times between 0700 – 1900 (Figure 1), since the natural photoperiod in southern Ontario fluctuates throughout the year and these hours encompass the minimum photoperiod experienced during this study.

**Figure 1:**
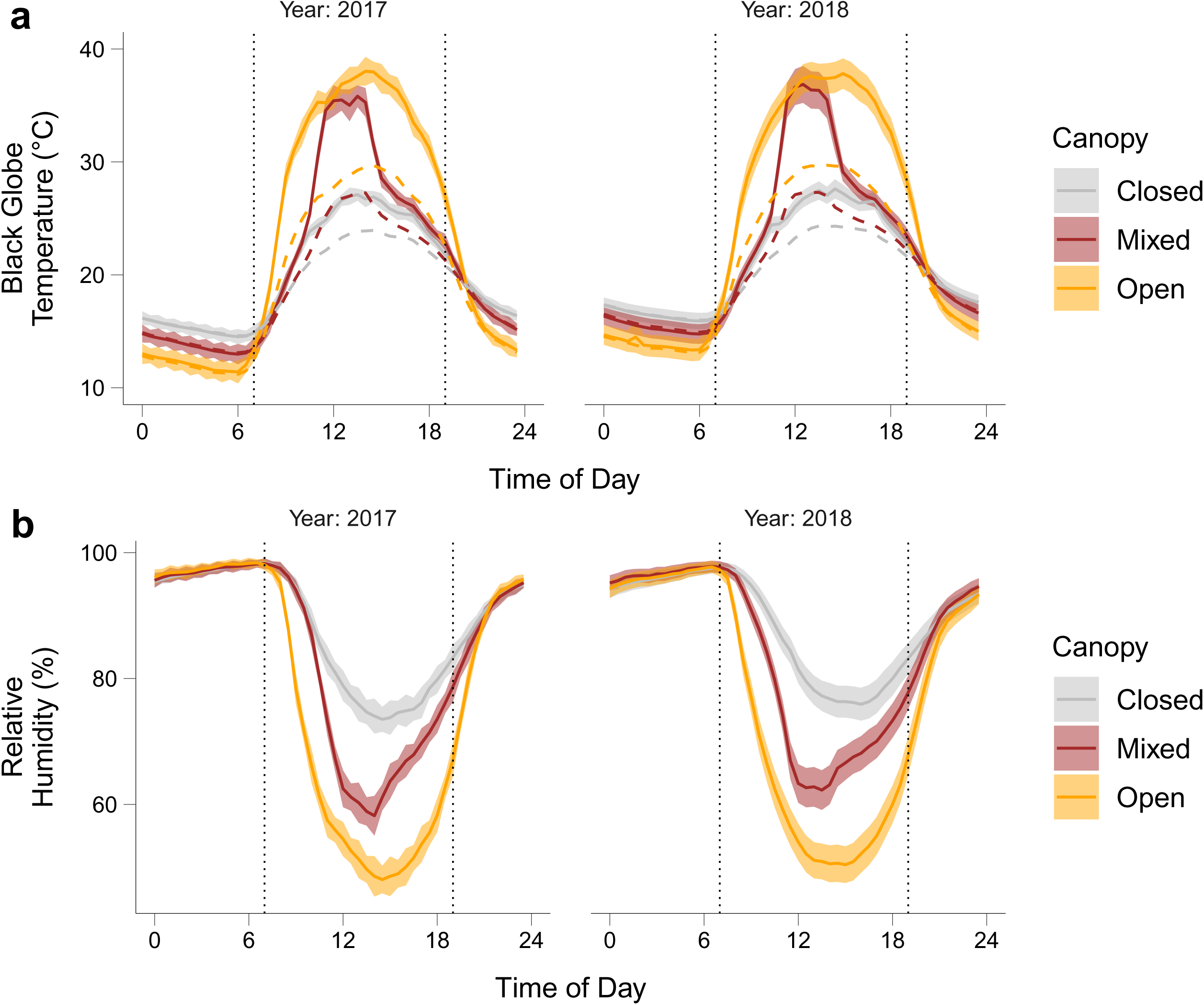
Averaged time of day trends of black globe temperature (a) and relative humidity (b) throughout the active season (May – September) in 2017 and 2018 across the three habitats of different canopy cover. Solid lines represent the mean, shaded areas indicate ±95% confidence interval (hourly). Dashed lines (a) represent the air temperature, for comparison. Black vertical dotted lines represent 0700 and 1900 and the area between them coincides approximately with the active period across the study period.

### T_e_ measurements

To capture the complexity of the thermal environment available to *T. s. sirtalis* within our study site we used T_e_ models. A series of copper snake models (Brock University Technical Services) designed to mimic the thermal properties of living snakes were deployed throughout the three canopy types (closed, mixed, and open) to record T_e_. Models were made entirely out of copper tubing with a soldered copper cap on one end, and a threaded, removable copper insert on the other. Copper has been used for several thermal biology studies as reptile analogs and has thermal transfer properties very similar to living snakes (Dzialowski 2005). Models work best when absorptivity, reflectance, size, and contact with substrate match their target species (Shine and Kearney 2001; Dzialowski 2005). Our models were manufactured to mimic scale colour and size of *T. s. sirtalis*, although they do exhibit large phenotypic variation in scale patterns and coloration within the study site. Therefore, models reflected an approximation of *T. s. sirtalis* inhabiting the study site. Estimates of juvenile *T. s. sirtalis*’s snout-vent length and body mass were taken and used to determine the dimensions of the models. Each model was 15.5 cm long, 2.5 cm in diameter (to accommodate iButton insertion), and filled with 30 mL of water to approximate the weight of a 30 g juvenile snake. Every model was sand-blasted and coated with non-reflective matte brown spray paint. Environmental variables such as humidity and wind speed may influence thermal properties of models, however, these typically result in negligible differences in overall mean temperatures (Shine and Kearney 2001).

Twenty-seven models were deployed across the field site, nine in each of the three distinct canopy types (closed, mixed, and open). Temperatures recorded by models reflect the T_b_ a living snake is likely to achieve in that microhabitat at that time in the absence of any regulatory behaviour. Ground-level microhabitat diversity available to snakes was incorporated by the placement of models within the three different habitats. The use of copper models bypassed two major obstacles: 1) wild *T. s. sirtalis* were not regularly disturbed, and 2) temperature measurements of multiple habitats could be simultaneously recorded. All models were fitted with temperature data loggers (Thermochron model DS1922L, Maxim Integrated™) to record T_e_ every 20 minutes for a total of 142 days each year from May 10^th^ to September 28^th^, 2017, and May 11^th^ to September 29^th^, 2018. Once the active season ended (after September), the dataset was assimilated into characteristic thermal profiles to make inferences about the thermal quality of each canopy type to *T. s sirtalis*.

### Model field placement

Models were placed in microhabitats accessible to snakes and to mimic realistic site selections of wild *T. s. sirtalis*. Both potential retreat and basking sites were selected. Models were placed beside and on top of coverboards, in woodpiles, in leaf litter, and beneath fallen logs. An approximately similar number of shaded and non-shaded microhabitats was chosen for all three habitats to avoid introducing biased model placements. For example, models placed on top of the boards were the most conspicuous and were also most likely to record the highest temperatures. Therefore, only two of the nine models from each habitat were placed on top of randomly selected coverboards. Each model had a unique numerical code and serial number, and each placement location was carefully marked so that individual models could be returned to their exact location after routine data downloads. Ground-level microhabitat diversity introduced by the placement of models was incorporated in the generation of generalized data of each habitat.

### Habitat d_e_

All snake models collected high-resolution temperature measurements every 20 minutes for 142 days through the reptile active season in Ontario (May – September). Over 500,000 temperatures were collected across the three habitats and two years. Although models collected temperatures continuously, our analysis was focused on temperature readings that occurred during a typical *T. s. sirtalis* active period at this latitude (∼ 43°N). *Thamnophis s. sirtalis* is strictly diurnal at this latitude, therefore, only temperatures recorded after sunrise and before sunset were used to analyze thermal quality (Figure 2). Data from all nine models within a habitat were used for analysis.

**Figure 2:**
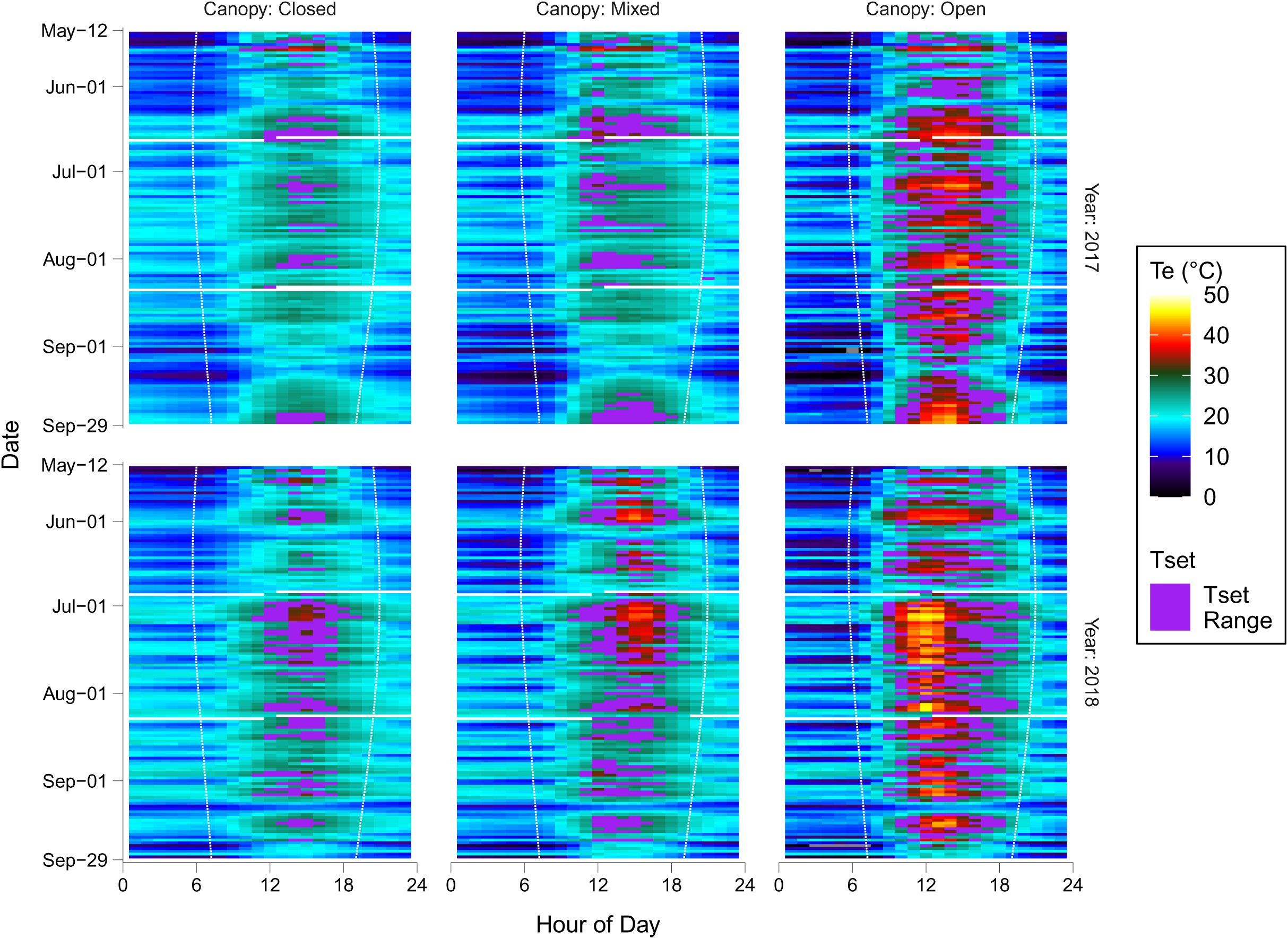
Tile plot of mean operative temperature (T_e_) retrieved from snake models from May – September 2017 and 2018 (with missing data depicted as white horizontal bars during iButton data retrieval and redeployment). Colour of each tile represents the mean value of all temperature readings at the respective site rounded to the nearest hour. Purple indicates blocks of time where T_e_ fell within the thermal preference (T_set_) range of *T. s. sirtalis*. White dashed lines represent sunrise (left) and sunset (right) for each individual day of the study period; the area within the white dashed lines coincides with the active period of *T. s. sirtalis*.

Habitat *d*_e_ is defined as the deviation from an animal’s T_set_ and is used to infer the thermal quality of a habitat (Hertz et al. 1993). Formally, the T_set_ is the range of selected temperatures between the 25^th^ (T_lower_) and 75^th^ (T_upper_) quartiles defining the temperature the animal would prefer under low-cost conditions (Hertz et al. 1993). Therefore, to estimate habitat *d*_e_ for all daytime T_e_ values recorded by the models throughout the active season, we calculated the deviation of T_e_ from the population-specific T_set_ range of *T. s. sirtalis* (T_lower_: 27.4°C, T_upper_: 32.1°C; (Giacometti et al. 2021) according to the following:

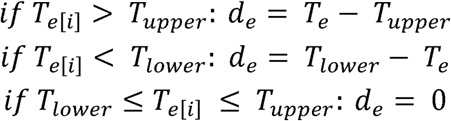

where T_e[i]_ represents an individual operative environmental temperature measurement taken from each *i*^th^ interval (20 minutes apart). T_e_ values which fall within the T_set_ bounds are assigned a *d*_e_ of 0, signifying optimal thermal quality. Mean *d*_e_ values could then be used to score each habitat in terms of thermal quality.

However, *d*_e_ may be limiting as it is highly right skewed (Figure S2), only represents the absolute deviation of T_e_ from T_set_, and assumes that deviations in temperatures above or below an animal’s thermal preference are equivalent (given they deviate by the same absolute value). This simplification can be hampering since the biological effects and response of being below the T_set_ range are vastly different than those of being above (Martin and Huey 2008); higher temperatures may induce heat stress (Gangloff and Telemeco 2018) whereas cooler temperature promote retreat site selection. To include the directionality of deviations from T_set_ we calculated the parameter, *d*_e_ raw, which is calculated similarly to *d*_e_ but will provide positive values if T_e_ is above T_set_ range and negative values if T_e_ is below T_set_ range, as follows:

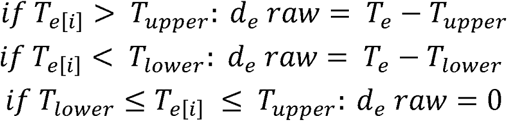

### Coverboard surveys

Coverboards were installed in the three habitats (closed, mixed, and open) in the winter of 2017, and we documented their occupation by snakes to determine habitat use of *T. s. sirtalis* throughout the active season. Nine coverboards were placed in each habitat using an aggregate design. Coverboards were arranged into aggregates of three boards 20 m from each other, with three aggregates in each habitat. Each aggregate was a minimum of 150 m away from all other aggregates and at least 400 m from the nearest road. All coverboards were identical 1.2 m x 2.4 m x 2 cm untreated plywood sheets. Coverboard surveys are a reliable method for documenting snake presence/absence that is safe and non-invasive and can be repeated for long study durations. Snakes can use coverboards as thermal and predation refugia as well as basking surfaces (Halliday & Blouin-Demers, 2015; Scheffers et al., 2009).

A total of 20 coverboard surveys (10 per year) were conducted throughout the reptilian active season in Ontario (May – September). *Thamnophis s. sirtalis* may leave hibernacula as early as March and remain active into October. We considered it more meaningful to focus on the species’ habitat use during the times of day and year with the greatest activity. We surveyed every habitat at approximately two-week intervals. Therefore, every coverboard was visited at least twice a month from May to September in 2017 and 2018. The sequence of habitat visitation was randomly chosen for a given survey period. Only one habitat was surveyed per excursion, although all 3 habitats were surveyed within days of each other to constitute a given survey number. All three aggregates within a habitat were surveyed within the same day. The sequence of aggregate visitation was randomly chosen. At the beginning of each survey, we recorded current weather conditions, approximation of cloud cover, and rainfall from the previous 24 hours from the nearest Environment Canada weather station (Port Colbourne, ON).

At each coverboard we recorded the start and end time of the coverboard visit, ambient temperature and RH using a handheld hygrometer (Extech Instruments™, model 445580). When snakes were encountered, the number of individuals, and location (beside the coverboard within 1 m, on top of the coverboard, or beneath the coverboard) were recorded. *T. s. sirtalis* were identified by three yellow longitudinal stripes (two lateral and one dorsal), a distinct head and lateral spotting (Giacometti et al. 2021). When *T. s. sirtalis* were encountered, they were immediately hand-caught and had morphometric measurements taken. We took snout-vent length (cm) and body mass (g) of every individual. Once all animals occupying the coverboard were vacated, we took the remaining environmental measurements; we took soil temperature readings using a probe thermometer (Extech Instruments™, model 39240, accuracy ± 1°C), and soil moisture readings using a soil moisture sensor (Delta-T™, model ML2x ThetaProbe, accuracy ±5%). For each measurement we took three spot measurements beneath the coverboard and recorded the mean value. Canopy cover photos were taken above each coverboard three times throughout the active season. Photos were processed in ImageJ to calculate average canopy cover for each habitat. Photos were taken with a wide-angle GoPro™ camera pointed directly towards the sky from the centre of the coverboard. Solar radiation (W/m^2^) was estimated at each coverboard using a pyranometer application (Hukseflux Thermal Sensors™) installed on a smartphone. The pyranometer application was calibrated using the Welland-Pelham weather station’s estimates of incoming solar radiation for that day.

### Incidental observations of T. s. sirtalis

Incidental sightings of *T. s. sirtalis* were compared among habitats to corroborate the habitat use obtained from coverboard encounters. During each survey we noted the total number of *T. s. sirtalis* we encountered while walking between aggregates within a habitat. These observations, referred to as incidental encounters, represent individuals that were actively basking or moving through the habitat. While walking between aggregates, a standardized search approach was employed. A snake hook was constantly swept through understory vegetation along the trail edges. We documented any snakes that were disturbed and positively identified. Most snakes were found basking directly on the trails or observed moving within a few meters of the trail edge. Snakes were identified by sight and were not captured or measured. *Thamnophis s. sirtalis* that were encountered in this manner were counted separately from those occupying coverboards. Incidental observations were standardized due to area searched and not the time spent *in-situ*. The same trails were always used and trail lengths for the closed forest (1.52 km), mixed shrub (1.54 km), and open peat (1.46 km) varied minimally (< 90 m). Variation in recorded survey time was mainly affected by time spent at coverboards and not the travel time between aggregates where incidental encounters happened meaning that survey duration was not appropriate to account for search effort.

### Skin surface temperatures of T. s. sirtalis in the field (T_b_)

We opportunistically measured skin surface temperatures (T_skin_) of active *T. s. sirtalis* observed in the field as a proxy for T_b_ using a thermal imaging camera (Spot Thermal Camera, model TG165 FLIR™) across both active seasons in 2017 and 2018. The opportunistic nature of these measurements led to inconsistent timing and locations of when measurements were recorded, since these individuals were not part of those observed during the active surveys. When imaged, snakes were actively basking or otherwise active within their environment and were a subset of the incidental observations of *T. s. sirtalis*. However, all measurements were captured from diurnally active snakes during the active season. Snakes found under coverboards were not imaged because they were intended for physical capture, which would have induced changes in temperature. Further, the time required for thermal image capture could have enabled their escape. However, some pilot thermal imaging was attempted on snakes under coverboards, which proved challenging given the high thermal homogeneity between the snakes and the ground. Since imaged snakes were not the ones we observed at coverboards, no specific habitat type, T_e_, or *d*_e_ can be ascribed to them.

### Data analysis

We performed all analyses using R (4.4.1) in RStudio (v. 2024.12.0.467) (R Core Team, 2024) with a significance level of 0.05. We used the ggplot2 (Wickham 2009) package to create our figures. To analyze effects of canopy type on thermal conditions, we fit linear mixed-effects models (LMMs) using the “lmer” function from the lme4 (Bates et al. 2015) package. We considered time T_e_ was within T_set_, time T_e_ was above T_set_, time T_e_ was below T_set_, and *d*_e_ raw as response variables, and we fit one model per response variable. In all models, we considered canopy type (categorical) and year (continuous) as fixed terms, and board number (categorical) as a random term. Fixed effect significances for LMMs were determined via Type III ANOVA using Satterthwaite’s method to estimate the degrees of freedom from the “anova” function in the lmerTest (Kuznetsova et al. 2017) package. To isolate specific canopy differences, we employed post-hoc tests using the emmeans (Lenth 2025) package, using the Tukey method p-value adjustment.

To analyze habitat use, we fit generalized linear mixed-effects models (GLMMs; family Poisson) using the “glmer” function from the lme4 (Bates et al. 2015) package. We considered *T. s. sirtalis* count at the coverboards and incidental count as response variables, fitting one model per response variable. For the *T. s. sirtalis* count model, we considered canopy type (categorical), *d*_e_ raw (continuous), and rainfall (continuous) as fixed terms, and survey number as a random term. For the incidental count model, we considered canopy type (categorical), survey length (continuous), and rainfall (continuous) as fixed terms, and survey number as a random term. Model fit was evaluated using the “check_model” function from the performance (Lüdecke et al. 2021) package. Analysis of deviance tables for GLMMs was performed via Type II Wald chi-square (χ^2^) test from the “Anova” function from the car (Fox and Weisberg 2019) package. To isolate specific canopy differences, we employed post-hoc tests using the emmeans (Lenth 2025) package, using the Tukey method p-value adjustment. We extracted partial effects for visualisation purposes (mean ± 95% CI) using the “ggeffect” function from the ggeffects (Lüdecke 2018) package.

## Results

### Habitat environmental characteristics

We observed large differences between T_a_ and T_bg_, even within the closed forest habitat (Figure 1). T_a_ and T_bg_ followed a similar daily pattern within each habitat type. Closed forest habitat was the coolest during the day/active period (0700 – 1900) but the warmest at night (1900 – 0700), thus being the most consistent thermal habitat. The open peat habitat was the warmest habitat during the day/active period (0700 – 1900) but the coolest at night (1900 – 0700) thus being the most extreme thermal habitat. RH was consistently high across all habitats at night. The closed forest had the highest daily RH while the open peat had the lowest. Mixed canopy habitat offered a shorter window of time for peak black globe temperatures, typically rising to high temperatures, similar to those seen in the open habitat, but only at midday, after which black globe temperatures in the mixed canopy converged toward those seen in the closed forest.

### T_e_ and d_e_

All habitats provided days where T_e_ was within the T_set_ range of *T. s. sirtalis* or *d*_e_ raw = 0 (Figure 2). T_set_ was not reached outside of the active period (sunrise – sunset) in any habitat. Across the 2 years of study the open habitat provided more days (242) where at any point T_e_ reached T_set_ than the mixed (149) and closed (100) habitats.

Canopy type (F_2, 24.654_ = 19.519, *p* < 0.001) and year (F_1, 25.371_ = 5.016, *p* = 0.034) both influenced the time T_e_ was within the T_set_ range of *T. s. sirtalis* (Table 2; 3). Based on post-hoc comparisons we observed a higher proportion of time T_e_ was within T_set_ in the open habitat than the mixed (*p* = 0.017) and closed (*p* < 0.001) habitats. The mixed habitat also exhibited a higher proportion of time T_e_ was within T_set_ than the closed habitat (*p* = 0.011). Both canopy type (F_2, 24.793_ = 27.602, *p* < 0.001) and year (F_1, 23.800_ = 16.504, *p* < 0.001) influenced the percentage of time T_e_ was below the T_set_ range of *T. s. sirtalis* (Table S1). The open habitat had a lower percentage of time when T_e_ was below T_set_ than the closed (*p* < 0.001) and mixed (*p* < 0.001) habitats (Table 3). There was no difference in the proportion of time T_e_ was below T_set_ between the closed and mixed habitats (*p* = 0.069). Canopy type (F_2, 24.531_ = 15.839, *p* < 0.001) and year (F_1, 23.411_ = 9.185, *p* = 0.006) both influenced the time T_e_ was above the T_set_ range of *T. s. sirtalis* (Table S2). The open habitat had a greater proportion of time when T_e_ was above T_set_ than the closed (*p* < 0.001) and mixed (*p* < 0.001) habitats (Table 3). There was no difference in the proportion of time T_e_ was above T_set_ between the closed and mixed habitats (*p* = 0.507) was found.

**Table 2:** Linear mixed effect model summary of time T_e_ was within T_set_ based on canopy and year.

| % Time $T_e$ Was in $T_{set}$ Range | | | | | |
| --- | --- | --- | --- | --- | --- |
| *Predictors* | *Estimates* | *CI* | *t value* | *p* | *df* |
| Canopy: Closed (Intercept) | 5.82 | 3.48 – 8.16 | 5.07 | <0.001 | 31.96 |
| Canopy: Mixed | 4.79 | 1.69 – 7.89 | 3.19 | 0.004 | 24.23 |
| Canopy: Open | 9.39 | 6.29 – 12.49 | 6.24 | <0.001 | 24.23 |
| Year: 2018 | 2.19 | 0.18 – 4.19 | 2.24 | 0.034 | 25.37 |
| **Random Effects** |  |  |  |  |  |
| $\sigma^2$ | 11.55 | | | | |
| $\tau_{00}$ Board | 3.94 | | | | |
| ICC | 0.25 |  |  |  |  |
| N Board | 27 |  |  |  |  |
| Observations | 50 |  |  |  |  |
| Marginal $R^2$ / Conditional $R^2$ | 0.509 / 0.634 | | | | |

**Table 3:** Summary table of proportion of time T_e_ was inside or outside T_set_ range (± SD) between sunrise and sunset. Superscripts indicate post-hoc comparisons across canopy type results (different letters indicate significant contrasts at α=0.05). Missing data from 2018 were from 2 open canopy boards (1.2 and 2.2) and in 2017 from 2 mixed canopy boards (2.3 and 3.1), due to iButton failure partway through study seasons.

|  | **Closed**<br>**(N=18)** | **Mixed**<br>**(N=16)** | **Open**<br>**(N=16)** |
| --- | --- | --- | --- |
| **% Time $T_e$ Was Within $T_{set}$ Range** | $6.91 \pm 3.80^a$ | $11.9 \pm 4.34^b$ | $16.2 \pm 4.00^c$ |
| **% Time $T_e$ Was Below $T_{set}$ Range** | $90.9 \pm 5.79^a$ | $82.1 \pm 8.89^a$ | $65.9 \pm 10.0^b$ |
| **% Time $T_e$ Was Above $T_{set}$ Range** | $2.21 \pm 2.77^a$ | $6.01 \pm 5.97^a$ | $17.9 \pm 10.3^b$ |

Both canopy type (F_2, 25.104_ = 13.431, *p* < 0.001) and year (F_1, 24.653_ = 9.860, *p* = 0.004) influenced *d*_e_ raw values (Table S3; Figure 3). We observed a greater *d*_e_ raw value in the open habitat than the mixed (*p* = 0.015) or closed (*p* < 0.001) habitats. We found no difference between the *d*_e_ raw values for the mixed and closed habitats (*p* = 0.119).

**Figure 3:**
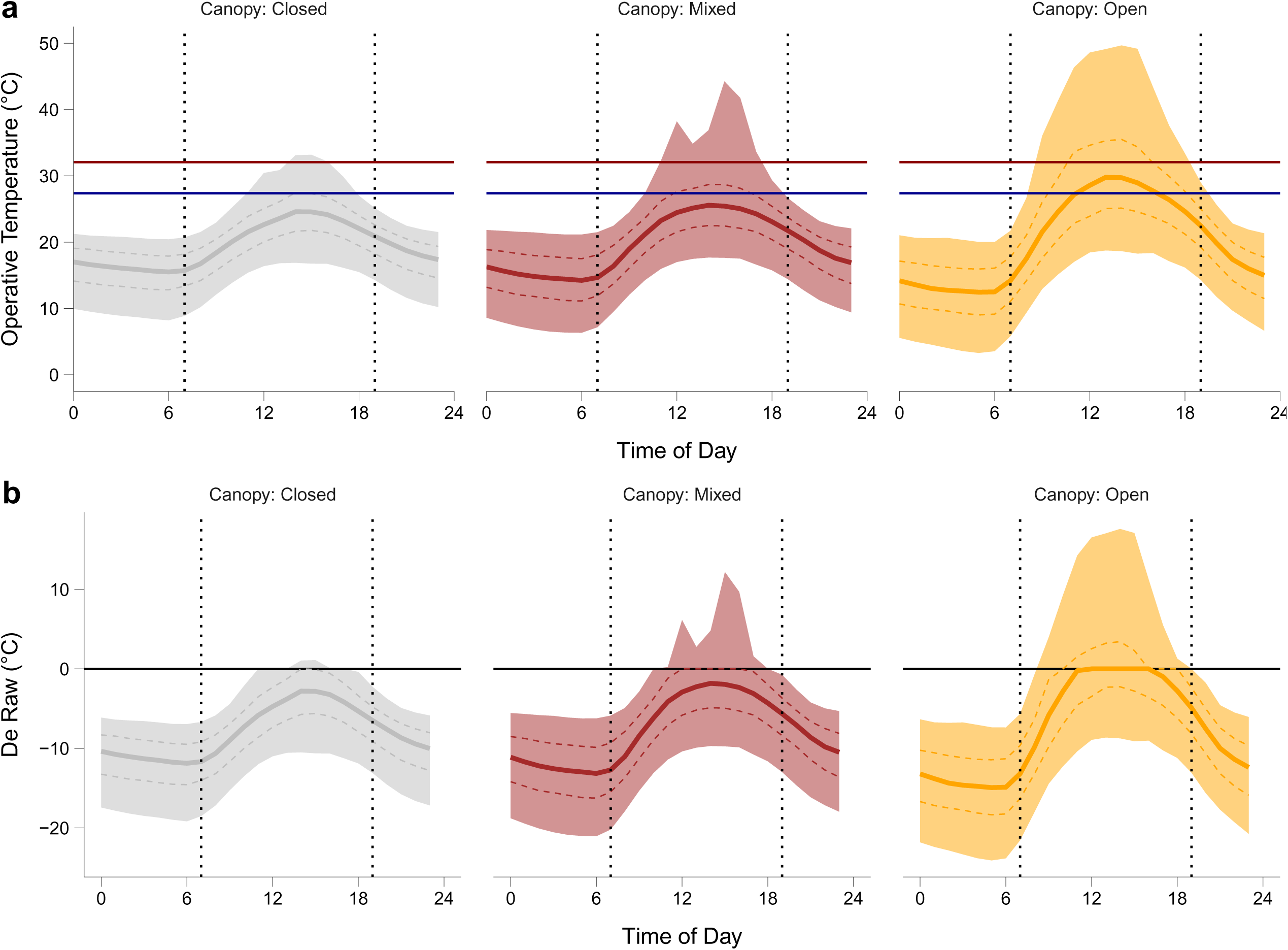
Time of day trends of environmental operative temperature (T_e_) (a) and d_e_ raw (b) throughout the active season (May – September) 2017 and 2018 across three canopy types. Black vertical dotted lines represent 0700 and 1900 and the area between them coincides with the active period within all days of the study. The blue and red horizontal solid lines (a) represents the lower (25^th^) bound and the upper (75^th^) bound of the thermal preference range of *T. s. sirtalis* respectively. Black horizontal solid line (b) represents a *d*_e_ raw value of 0 (i.e. when T_e_ falls within T_set_). Solid lines represent the mean, dashed lines 25^th^ and 75^th^ percentiles, and shaded area the 95% confidence interval of T_e_ (a) and *d*_e_ raw (b). Values correspond to the mean values across all models and across all days of the study period (May – September 2017 and 2018) rounded to the nearest half hour.

### Habitat use

Based on coverboard surveys, both canopy type (χ^2^_df=2_ = 10.942, *p* = 0.004) and *d*_e_ raw (χ^2^_df=1_ = 5.822, *p* = 0.016) influenced *T. s. sirtalis* count (Table 4; Figure 4). Rainfall had a minor effect on *T. s. sirtalis* count (χ^2^ = 3.705, *p* = 0.054; Table 4). Based on post-hoc comparisons we observed a higher use of the mixed shrub habitat compared to either the closed forest (*p* = 0.026) or open peat habitat (*p* = 0.037). No difference in *T. s. sirtalis* count was found between the closed forest and open peat habitats (*p* = 0.960). We observed a total of 27 and 31 snakes in 2017 and 2018, respectively; 91% of snakes we observed were under coverboards.

**Figure 4:**
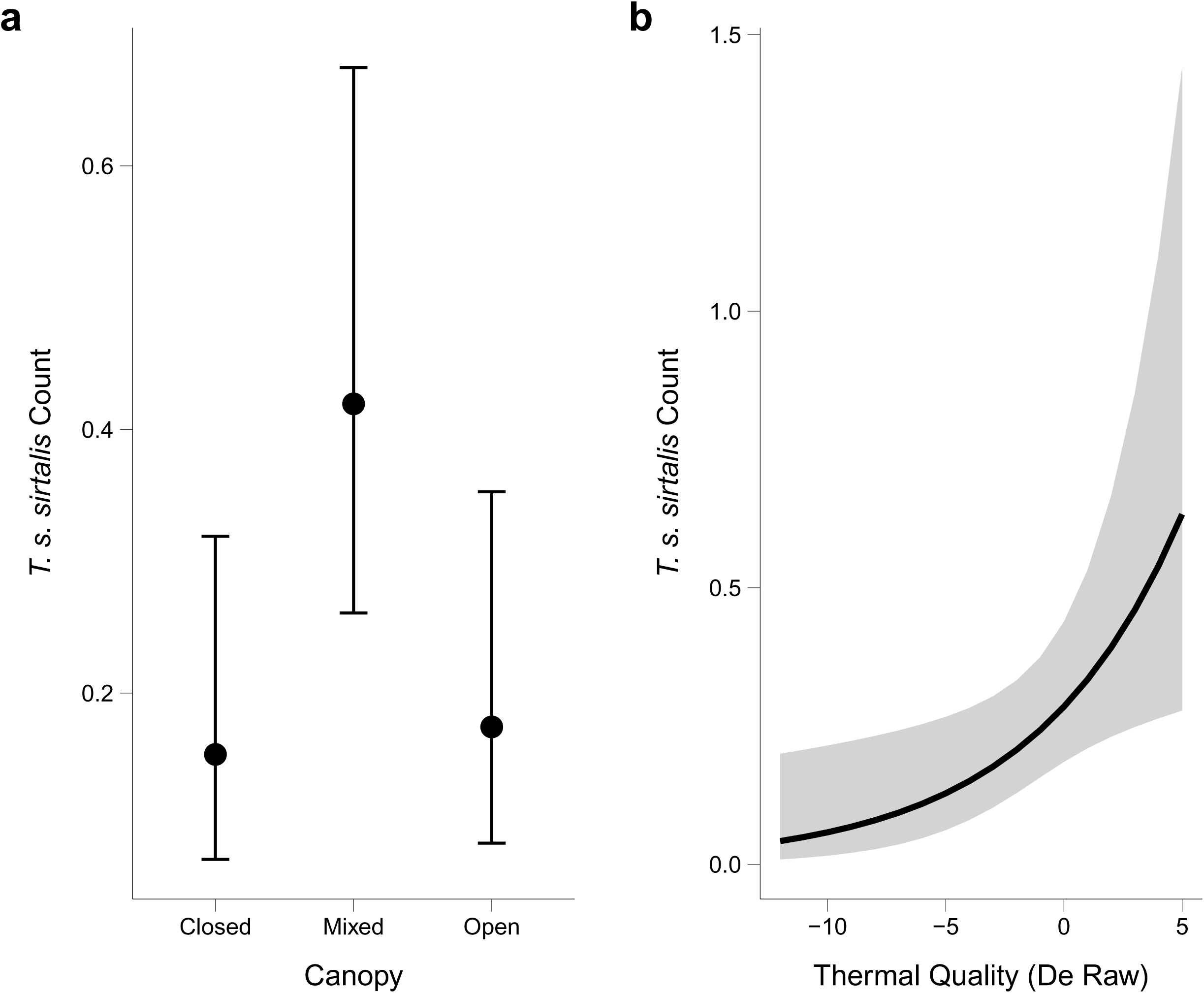
Summary model fits (estimate ± 95% confidence interval) depicting how *T. s. sirtalis* count per aggregate is influenced by habitat (a) and thermal quality (b).

**Table 4:** Generalized linear mixed effects model of *T. s. sirtalis* count from coverboard surveys based on canopy, *d*_e_ raw, and rainfall with survey number as a random effect.

| *T. s. sirtalis* |  |  |  |  |
| --- | --- | --- | --- | --- |
| *Predictors* | *Incidence Rate Ratios* | *CI* | *F-stat* | *p* |
| Canopy: Closed (Intercept) | 0.15 | 0.07 – 0.32 | -5.02 | **<0.001** |
| Canopy: Mixed | 2.73 | 1.28 – 5.84 | 2.59 | **0.010** |
| Canopy: Open | 1.14 | 0.44 – 2.91 | 0.27 | 0.790 |
| DeRaw | 1.73 | 1.11 – 2.71 | 2.41 | **0.016** |
| Rainfall | 1.26 | 1.00 – 1.59 | 1.92 | 0.054 |
| **Random Effects** |  |  |  |  |
| $\sigma^2$ | 1.40 | | | |
| $\tau_{00}$ Survey | 0.28 | | | |
| ICC | 0.16 |  |  |  |
| N<sub>Survey</sub> | 20 |  |  |  |
| Observations | 180 |  |  |  |
| Marginal R<sup>2</sup> / Conditional R<sup>2</sup> | 0.238 / 0.363 |  |  |  |

Canopy type (χ^2^_df= 2_ = 12.944, *p* = 0.002) affected incidental *T. s. sirtalis* count (Table 5). There was a difference between the mixed and open canopies (*p* = 0.001) with fewer in the open canopy. There was no effect of rainfall on incidental *T. s. sirtalis* count (χ^2^_df=1_ = 0.310, *p* = 0.578).

**Table 5:**
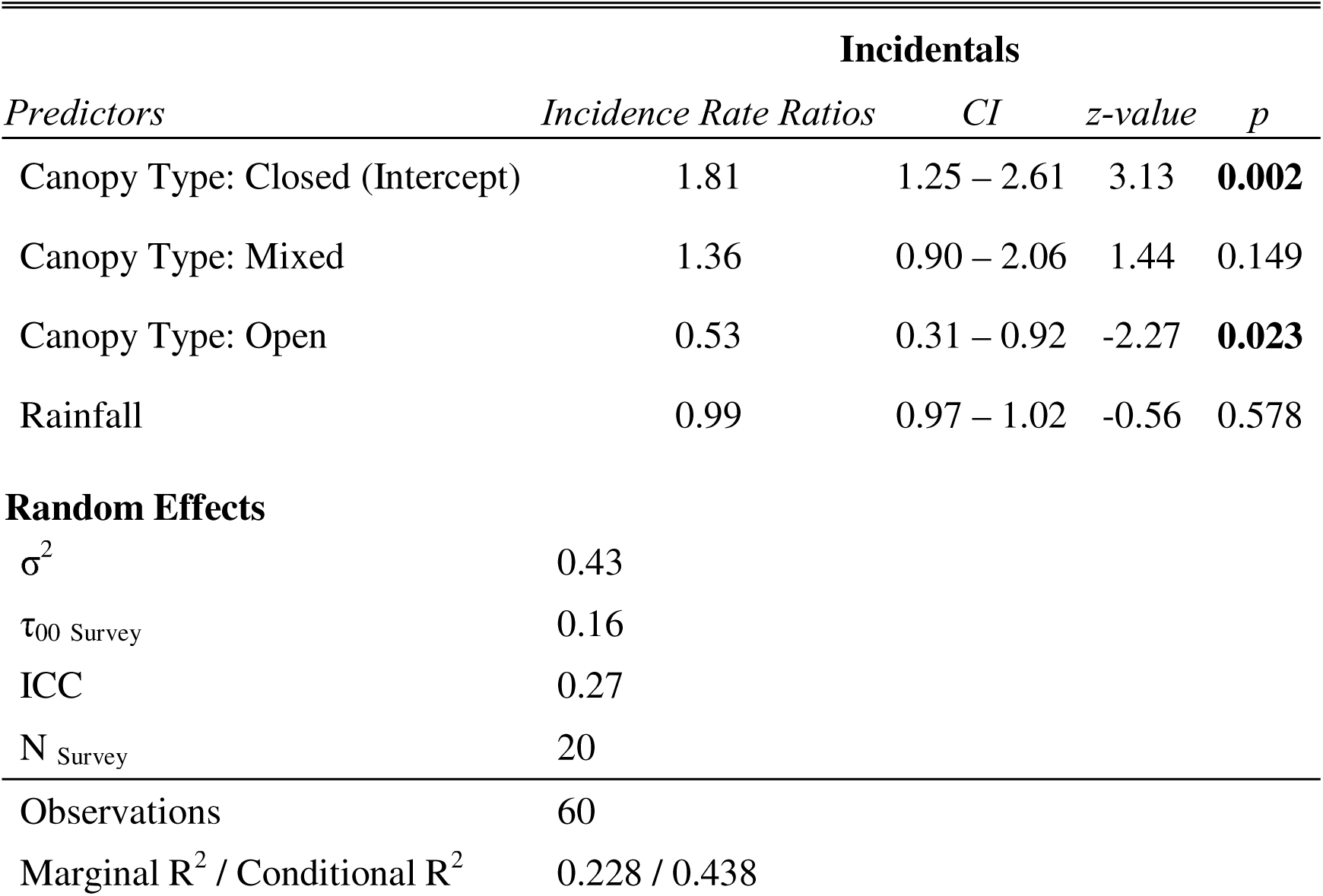
Generalized linear mixed effects model of incidental *T. s. sirtalis* count during surveys based on canopy type and survey duration.

### T. s. sirtalis skin surface temperature in the field

During the active seasons (May – September) of 2017 and 2018 the mean T_b_ of *T. s. sirtalis* was 28.1 ± 1.7 °C. This mean skin surface temperature is within *T. s. sirtalis*’s thermal preference range (T_set_; 27.4 °C – 32.1 °C) (Giacometti et al. 2021). Nearly two thirds (29 of 44, 65.9%) of active snakes had T_b_ within the T_set_ range of *T. s. sirtalis*. The remaining snakes (15 of 44, 34.1%) had T_b_ below *T. s. sirtalis*’s thermal preference range, ranging from <0.1 – 2.5 °C below the T_set_ range. No snake had a T_b_ above *T. s. sirtalis*’s thermal preference range.

## Discussion

In this study we set out to uncover how the environmental thermal profile influences habitat selection in *T. s. sirtalis*. We determined that the closed-forest, mixed-shrub, and open-peat habitats all provided unique thermal habitats. Based on precepts of thermal ecology, we hypothesized that the habitat with the highest thermal quality (lowest investment into thermoregulation needed to obtain a T_b_ within T_set_ range) would be the most used by *T. s. sirtalis* (Row and Blouin-Demers 2006a; Halliday and Blouin-Demers 2016). The open canopy provided the warmest temperatures and the best thermal quality for *T. s. sirtalis* throughout their active season. However, *T. s. sirtalis* were the least abundant in the open canopy for both coverboard (not different than closed canopy) and incidental encounters. The greatest abundance of *T. s. sirtalis* occurred in the mixed canopy for both coverboard and incidental encounters (not different than closed canopy). This finding may indicate that thermoregulation is not the most important factor influencing *T. s. sirtalis* habitat use or that thermal quality oversimplifies the thermal landscape and does not capture all thermal aspects of a habitat. We also hypothesized that *T. s. sirtalis* would be the most abundant at *d*_e_ raw values near zero (Hertz et al. 1993; Row and Blouin-Demers 2006b). Coverboard count increased as *d*_e_ raw increased even beyond zero (highest thermal quality) meaning that we found more snakes at temperatures above their T_set_ range than we did at temperatures within their T_set_ range. The disconnect between finding more snakes at higher temperatures and finding the least number of snakes in the hottest habitat may be due to snakes occupying the mixed canopy habitat more than the open canopy habitat but using coverboards as thermal refuges. The use of coverboards as thermal refuges would explain why we found more snakes at high temperatures as they are trying to prevent themselves from overheating by remaining stationary in a cooler microhabitat.

### The Goldilocks effect

The tale of Goldilocks embodies the idea that when provided with two opposing options a third intermediate option is preferred. We found that *T. s. sirtalis*’s habitat use in respect to temperature follows a Goldilocks effect; the closed-forest habitat was too cold, the open-peat habitat was too hot, but the mixed-shrub habitat was just right. For temperate snakes, warmer habitats are preferred as temperature can be their most limiting resource (Row and Blouin-Demers 2006a, 2006b; Halliday and Blouin-Demers 2016). If temperature is the main factor influencing the habitat usage of *T. s. sirtalis* then the habitat that provides the highest thermal quality (open canopy) should be preferred. We found that the mixed canopy habitat was favoured by *T. s. sirtalis* despite having a lower thermal quality than the open canopy habitat. The mixed canopy habitat might be more desirable to *T. s. sirtalis* due to thermal heterogeneity, resource abundance (e.g. food and retreat sites), and predation risk.

The mixed canopy habitat potentially provided the best mix of cover and open basking areas for active thermoregulation and predator avoidance. Snakes prefer heterogenous habitats that allow them to perform thermoregulatory behaviours for both cooling and warming (Blouin Demers and Weatherhead 2002). This thermal heterogeneity is present in the mixed canopy habitat with sunny areas for basking and retreat sites for cooling. Stationary models fail to capture the spatial thermal heterogeneity of a habitat and thus thermal heterogeneity is not expressed in thermal quality values. Without input from spatial thermal heterogeneity (Llanos-Garrido et al. 2023), thermal quality may be an oversimplification of the thermal landscape and may not accurately predict the habitat use of temperate snakes (Sears et al. 2016).

### The highest thermal quality is not always the best

We anticipated the habitat with the highest thermal quality (*d*_e_ raw closest to zero) would have the greatest abundance of snakes as they would have to invest the least amount of energy into thermoregulation (Hertz et al. 1993; Row and Blouin-Demers 2006a, 2006b; Thompson et al. 2018). The open canopy had the best average thermal quality across the study and the highest proportion of time T_e_ was within T_set_. Therefore, we had predicted the open habitat would be the most used habitat by *T. s. sirtalis*, as it is with other temperate snake species (Row and Blouin-Demers 2006a, 2006b; Halliday and Blouin-Demers 2016; Diaz and Blouin-Demers 2017). However, the open canopy had the lowest *T. s. sirtalis* count for both coverboard and incidental encounters, which may simply be related to heat stress. Due to the open canopy’s warmer temperatures, the open canopy habitat experienced the most time T_e_ was above T_set_ along with the most extremely high temperatures. Ectotherm’s thermal performance curves are nonlinear and asymmetric with a much steeper decline in performance above T_set_ than below (Huey and Slatkin 1976; Van Berkum 1988; Martin and Huey 2008; Huey and Pianka 2017). A steeper decline in performance above T_set_ may lead ectotherms to trade-off thermal optimality with thermal tolerance and thus explain the selection of thermally suboptimal habitats (Martin and Huey 2008).

The high T_e_ variation in the open canopy habitat shows how T_e_ variability may be an overlooked parameter of *d*_e_. A fundamental flaw of *d*_e_ is that it oversimplifies complex and variable thermal landscapes (Vickers et al. 2011). Ectotherms are not perfect thermoregulators and experience a range of body temperatures when in thermally fluctuating environments (Martin and Huey 2008; Giacometti et al. 2023). Temperate snakes try to maintain a more stable T_b_ by using thermally stable retreat sites or refugia (Blouin Demers and Weatherhead 2002; Hecnar and Hecnar 2011; Lorioux et al. 2013; Cox et al. 2018). In our study site, the open habitat is a previously mined wetland that consequently has few thermal refugia (Yagi and Litzgus 2012). The lack of thermal refugia likely restricts *T. s. sirtalis*’s ability to inhabit the open canopy habitat as they cannot efficiently take shelter from high temperatures (Webb and Shine 1998). This finding corroborates the prediction that since reptiles do not have a physiological thermal-safety margin, as T_e_ often exceeds their CT_max_ (Sunday et al. 2014), then behaviour must be the primary means of overheating avoidance (Sinervo et al. 2024). In the current study, despite the open canopy habitat providing the lowest *d*_e_ values, *T. s. sirtalis* limits its use of this habitat as they possibly lack a way of physiologically buffering against high temperatures. Alternatively, habitat use in *T. s. sirtalis* may be influenced by factors beyond temperature, which may have had a larger impact than we anticipated. The open canopy habitat may be the least preferred despite having the highest thermal quality due to increased predation risk from aerial predators (Webb and Whiting 2005; Duchesne et al. 2022) or milk snakes (Row and Blouin-Demers 2006a, 2006b) which frequent open canopy habitats. Overall, thermal quality alone may not be the most important factor influencing *T. s. sirtalis* habitat use; thermal stability, and avoidance of extreme temperatures and predators may also play a role.

### Higher temperatures mean more snakes

Despite the warmest habitat not having the greatest abundance of snakes, we found more *T. s. sirtalis* under coverboards at higher *d*_e_ raw values. This finding is contrary to our hypothesis that snakes would be found in the greatest numbers when *d*_e_ raw was equal to or just below zero (Hertz et al. 1993; Martin and Huey 2008). More snakes using coverboards at high temperatures could be a function of them using the coverboards as thermal retreat sites. Coverboards provide a more consistent temperature than the surrounding air and shade the substrate beneath them from direct solar radiation (Engelstoft and Ovaska 2000; Eye et al. 2018). Therefore, snakes may use coverboards to remain warm overnight or to retreat from extreme heat during the day. If snakes are using coverboards to escape the temperatures of their surrounding habitat, they would not be a useful method of assessing habitat usage by active snakes. However, a greater abundance of snakes under coverboards, even if inactive, is still an indicator of a greater number of snakes using that habitat. Indeed, inactive snakes under coverboards may simply be taking advantage of warm refugia to aid in digestion, a process that can last between 2 to 6 days (Stevenson et al. 1985; Bessler et al. 2010).

A potential challenge with our survey is that we could not assign incidental encounters a T_e_ value as they were not found near a model, meaning we cannot analyse how thermal quality impacts this type of use of the habitat. These incidental encounters represent snakes that are actively moving around the habitat and are the best indicators of habitat use. Another shortcoming is that all the coverboard surveyed snakes were found underneath the coverboards, but the models used to predict their temperature were not all placed beneath the coverboards. Therefore, even the snakes found at coverboards which we can assign T_e_ to may not be experiencing the same temperatures as the model.

### Possible limitations

There were a few limitations with this study’s experimental design. First, we had low encounter rates which may be due to our coverboards. We used new coverboards set in novel locations within the first year of the study. Coverboards take time to be discovered by snakes and may require several months to become sites snakes frequent (Engelstoft and Ovaska 2000; Scheffers et al. 2009), although we did encounter similar total numbers across both years. Another complication with our study is whether coverboards reflect habitat use or thermal refuge use. Since the substrate underneath a coverboard is not exposed to solar radiation it can remain at cooler temperatures than the substrate that makes up the rest of the habitat. At high temperatures, snakes we find under coverboards may be using the coverboard to cool down. Therefore, the abundance of snakes we find at coverboards will only reflect snakes in a particular habitat seeking thermal refuge and reflect the total number using the habitat. *Thamnophis sirtalis* using coverboards as thermal refugia would explain why we found more snakes at high *d*_e_ raw values. At positive *d*_e_ raw values we may have found more snakes at coverboards because they were seeking a thermal refuge from the surrounding habitat which was at temperatures above their T_set_ range. Furthermore, since our models were not all placed beneath coverboards, but all snakes found at coverboards were beneath them, our T_e_ and *d*_e_ raw values do not reflect the exact temperatures these snakes were experiencing. Therefore, we may be overestimating the T_e_ and *d*_e_ values experienced within each habitat and the correlation of these values with *T. s. sirtalis* abundance.

Finally, we worked under the assumption that temperature was the main determinant of habitat use in temperate snakes (Row and Blouin-Demers 2006a, 2006b; Lelièvre et al. 2011; Lorioux et al. 2013; Halliday and Blouin-Demers 2016). In nature, however, there are many factors that impact habitat use such as resource availability (Madsen and Shine 1996), predator avoidance (Webb and Whiting 2005), interactions (both competitive and mutually beneficial) with conspecifics (Halliday and Blouin-Demers 2017a), and even idiosyncratic (i.e. local) habitat characteristics (Pafilis et al. 2019). Thus, our findings of *T. s. sirtalis* using the mixed canopy more than the closed or open canopy could be related to factors outside of the T_e_ provided by each habitat. While the mixed canopy may have provided the best mix of cover and open basking areas for active thermoregulation and predator avoidance (Webb and Whiting 2005; Row and Blouin-Demers 2006a, 2006b; Duchesne et al. 2022), there may have also been more soft bodied invertebrate prey within the damper and shadier, mixed and closed canopies. Evaluating these alternative hypotheses were beyond the scope of the current study but nonetheless warrant further investigation to understand which fine scale factors inform habitat use in *T*. *s*. *sirtalis*.

### Skin surface temperature reaffirms T_set_

In small to medium sized snakes T_skin_ can be used as a proxy for T_b_ due to their high correlation (Halliday & Blouin-Demers, 2017). Therefore, in *T. s. sirtalis* we can use T_skin_ to estimate if a snakes T_b_ is within T_set_. The mean T_skin_ of the 44 wild snakes we measured was 28.1°C, falling within but closer to the low end of *T. s. sirtalis*’s T_set_ range (27.4 °C – 32.1 °C) (Giacometti et al. 2021). Most of the snakes we measured were within the T_set_ range with the remaining being below the T_set_ range. This distribution of T_skin_ being within or below T_set_ is expected. In general, the T_set_ may be an overestimation of the T_b_ range ectotherms strive to maintain in the wild (Martin and Huey 2008), since there are costs associated with thermoregulation, such as increased predation risk. Therefore, temperate ectotherms may benefit from maintaining a slightly lower T_b_ rather than incur the costs of thermoregulation. Furthermore, our T_skin_ values may be lower based on the snakes we sampled, since these snakes were stationary and not under a coverboard. Since *T. s. sirtalis* is a highly active species, most stationary individuals that are out in the open are likely to be basking. Basking snakes would have lower body temperatures as they are actively trying to increase their T_b_.

### Conclusions

Our findings contrast the hypothesis that snakes seek out habitats that offer the greatest thermal quality (Hertz et al. 1993). We found that *T. s. sirtalis* prefers an intermediate/mixed habitat that provides a less limiting thermal landscape than a closed canopy habitat but is sheltered from the thermal extremes of an open canopy habitat. From these findings we believe that avoiding temperatures above T_set_ may be more important than thermal quality. Given that the physiological consequences of overheating likely influence ectotherms to avoid achieving temperatures above their T_set_, it is important to factor in the directionality of temperature deviation when calculating thermal quality. Our findings also contradict previous findings that temperate snakes near the northern extent of their range prefer the warmer open habitats (Row and Blouin-Demers 2006a, 2006b; Halliday and Blouin-Demers 2016; Diaz and Blouin-Demers 2017). Overall, habitat use of *T. s. sirtalis* may follow a Goldilocks principle: habitats that are too hot (open canopy) or too cold (closed canopy) are used less than habitats that are just right (mixed canopy). However, due to limitations our coverboard encounter rates may poorly reflect habitat use of the species, and our T_e_ values may be an overestimation of temperatures experienced by survived snakes. The correlation of more *T. s. sirtalis* found under coverboards with high *d*_e_ raw values may indicate that snakes are using coverboards as thermal refuges. Therefore, coverboards may not be good indicators of habitat use for thermal studies as they represent thermal refugia, especially in relatively active diurnal snakes such as *T. sirtalis* (Larsen 1987; Charland and Gregory 1995). Future work is needed to investigate the use of coverboards as thermal retreat sites by snakes to assess coverboard survey reliability as a means of quantifying snake habitat use.

## Declarations

Conflicts of interest

None

## Ethics and permit approval

Brock AUP Protocol #17-01-01 and MNRF Wildlife Scientific Collectors Authorisation #1086633

## Availability of data and code

Data are available in the Borealis data repository: https://doi.org/10.5683/SP3/UVAQ7M

## Acknowledgements

We would like to thank to Theresa Bukovics, Byron Garcia, and Alanah Joyce who helped find snakes during survey work for 8Trees Environmental Consultants. We also thank Anne Yagi for introducing us to the field site and her assistance with logistics. Research funding was provided by a Natural Sciences and Engineering Research Council of Canada Discovery Grant to GJT (RGPIN-2014-05814).

## Data Availability Statement

Data are available at the following data repository: https://doi.org/10.5683/SP3/UVAQ7M

