## Supplementary Figure 1 for "Operative temperatures of eastern garter snakes (*Thamnophis sirtalis sirtalis*) reveal a Goldilocks effect for habitat use"

Figure S1: Tile plots of mean relative humidity (RH; a) and air temperature (T_a_; b) from May – September 2017 and 2018 (with missing data depicted as white horizontal bars during iButton data retrieval and redeployment). Colour of each tile represents the mean value of RH (a) or T_a_ (b) readings rounded to the nearest hour. White dashed lines represent sunrise (left) and sunset (right) for each individual day. Area within the white dashed lines coincides with the active period of *T. s. sirtalis*.
