## Supplementary Figure 2 for "Operative temperatures of eastern garter snakes (*Thamnophis sirtalis sirtalis*) reveal a Goldilocks effect for habitat use"

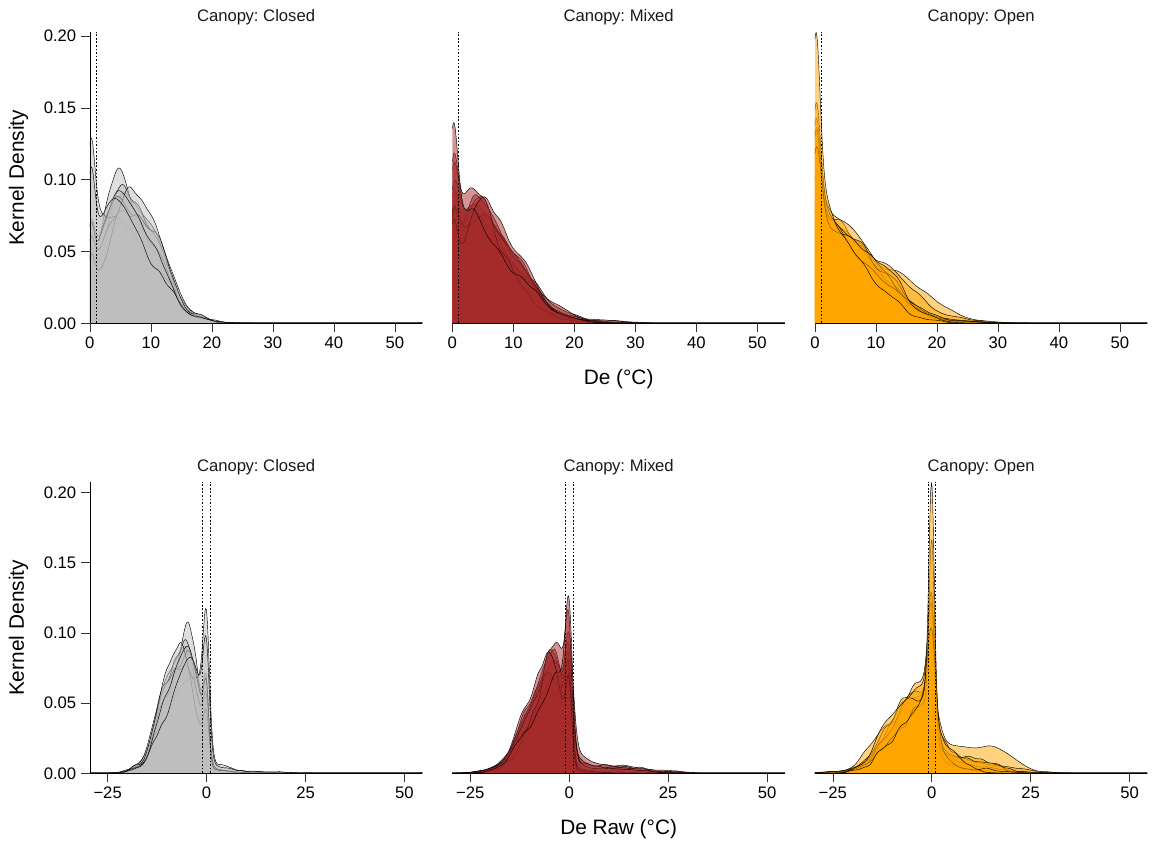

Figure S2: Kernel density plots of *d*_e_ (a) and *d*_e_ raw (b) based on canopy type. Horizontal dashed lines represent *d*_e_ (a) and *d*_e_ raw (b) values of 1 and -1 (b only).
