## Supplementary Table 1 for "Operative temperatures of eastern garter snakes (*Thamnophis sirtalis sirtalis*) reveal a Goldilocks effect for habitat use"

Table S1: Linear mixed effect model summary of time T_e_ was below T_set_ based on canopy and year.

| **% Time T_e_ Was Below T_set_ Range** | | | | |  |
| --- | --- | --- | --- | --- | --- |
| *Predictors* | *Estimates* | *CI* | *t value* | *p* | *df* |
| Canopy Type: Closed (Intercept) | 93.55 | 88.38 – 98.73 | 37.10 | **<0.001** | 27.04 |
| Canopy Type: Mixed | -8.14 | -15.31 – -0.98 | -2.34 | **0.028** | 24.52 |
| Canopy: Open | -25.37 | -32.54 – -18.20 | -7.30 | **<0.001** | 24.52 |
| Year: 2018 | -5.34 | -8.06 – -2.63 | -4.06 | **<0.001** | 23.80 |
| **Random Effects** | | | | | |
| σ^2^ | 20.35 | | | | |
| τ_00_ _Board_ | 43.17 | | | | |
| ICC | 0.68 | | | | |
| N _Board_ | 27 | | | | |
| Observations | 50 | | | | |
| Marginal R^2^ / Conditional R^2^ | 0.650 / 0.888 | | | | |
