## Supplementary Table 2 for "Operative temperatures of eastern garter snakes (*Thamnophis sirtalis sirtalis*) reveal a Goldilocks effect for habitat use"

Table S2: Linear mixed effect model summary of time T_e_ was above T_set_ based on canopy and year.

| **% Time T_e_ Was Above T_set_ Range** | | | | |  |
| --- | --- | --- | --- | --- | --- |
| *Predictors* | *Estimates* | *CI* | *t value* | *p* | *df* |
| Canopy Type: Closed (Intercept) | 0.59 | -3.87 – 5.06 | 0.27 | 0.787 | 26.48 |
| Canopy Type: Mixed | 3.39 | -2.81 – 9.60 | 1.13 | 0.270 | 24.28 |
| Canopy: Open | 16.12 | 9.92 – 22.33 | 5.36 | **<0.001** | 24.28 |
| Year: 2018 | 3.23 | 1.03 – 5.43 | 3.03 | **0.006** | 23.41 |
| **Random Effects** | | | | | |
| σ^2^ | 13.30 | | | | |
| τ_00_ _Board_ | 33.38 | | | | |
| ICC | 0.72 | | | | |
| N _Board_ | 27 | | | | |
| Observations | 50 | | | | |
| Marginal R^2^ / Conditional R^2^ | 0.516 / 0.862 | | | | |
