## Supplementary Table 3 for "Operative temperatures of eastern garter snakes (*Thamnophis sirtalis sirtalis*) reveal a Goldilocks effect for habitat use"

Table S3: Linear mixed effect model summary of *d*_e_ raw based on canopy and year.

|  | **Mean *d*_e_ raw** | | | | |
| --- | --- | --- | --- | --- | --- |
| *Predictors* | *Estimates* | *CI* | *t value* | *p* | *df* |
| Canopy Type: Closed (Intercept) | -7.05 | -7.92 – -6.19 | -16.68 | **<0.001** | 28.74 |
| Canopy Type: Mixed | 1.19 | 0.00 – 2.37 | 2.07 | **0.049** | 24.75 |
| Canopy: Open | 2.96 | 1.78 – 4.15 | 5.16 | **<0.001** | 24.75 |
| Year: 2018 | 0.85 | 0.29 – 1.41 | 3.14 | **0.004** | 24.65 |
| **Random Effects** | | | | | |
| σ^2^ | 0.87 | | | | |
| τ_00_ _Board_ | 1.01 | | | | |
| ICC | 0.54 | | | | |
| N _Board_ | 27 | | | | |
| Observations | 50 | | | | |
| Marginal R^2^ / Conditional R^2^ | 0.468 / 0.753 | | | | |
